# Evolutionary Histories and Environments Shape Ugandan and Global Oral Microbiomes

**DOI:** 10.64898/2026.05.20.726600

**Authors:** Iyunoluwa J. Ademola-Popoola, Kathleen Grogen, Muslihudeen A. Abdul-Aziz, K. Ta Christine, Karen Tang, Ran Blekham, Luis B. Barreiro, George Perry, Laura S. Weyrich

## Abstract

Industrialization has been identified as the single biggest factor driving global microbiome diversity. While many studies examining gut microbiomes attribute these shifts to dietary increases in fat and reductions in protein, oral microbiome responses to industrialization remains debated. The oral microbiome is more resilient due to long-standing coevolution with host tissues and biofilm stability. However, limited geographic and historical representation has constrained our understanding of how these transitions unfolded globally in the oral microbiomes. Here, we investigate oral microbiome variation in Batwa rainforest hunter-gatherers and neighboring Bakiga subsistence farmers from southwestern Uganda, comparing them with publicly available data from Tanzanian, Venezuelan, and industrialized populations from North America, Europe, and Australia. Using 16S rRNA gene sequencing, we characterized salivary microbiota and evaluated differences in local and global diversity, composition, and differential abundance. Ugandan populations contained significant compositional differences but similar levels of diversity, suggesting that shared environments and dietary overlap may shape microbial assemblages despite distinct cultural histories. Globally, strong continental and industrialization effects were observed in the oral microbiome, with all industrial populations clustering separately from people living in other locations. African populations also clustered separately from non-African groups. Oral microbiome diversity was highest in Ugandan individuals and lowest in industrialized populations, mirroring patterns previously observed in the gut microbiome. Together, these findings demonstrate that both geography and subsistence strategy structure global oral microbiome variation. They also clarify the position that oral microbial communities record biocultural transitions and highlight the need to better understand the industrial mechanisms that shape microbial diversity in the oral cavity.

## INTRODUCTION

Factors that shape modern gut microbiome diversity, including diet, migration, and antibiotic usage, are increasingly well described on a global scale, offering key insights into how microbial communities in distinct human populations are structured across time and space (David et al., 2014; De Filippo et al., 2010; Rosas-Plaza et al., 2022; Schwartz et al., 2020; Singh et al., 2017; van der Vossen et al., 2023; Voreades et al., 2014; Yassour et al., 2016). However, the large-scale evolutionary, social, and cultural factors that drive oral microbiome variation therefore remain poorly understood. Unlike the copious studies in the gut (reviewed by Davenport et al., 2017), how the composition and structure of the oral microbiome have changed in response to evolutionary-scale forces remains is more limited (De Filippis et al., 2014; Hansen et al., 2018; Lassalle et al., 2018). This gap is complicated by the inherent complexity of microniches within the oral cavity, as well as inconsistencies across studies that use different populations, sequencing approaches, and analytical pipelines.

Recent research has now explored how factors such as industrialization—one of the most significant drivers of global gut microbiome diversity (Quercia et al., 2014; Sonnenburg & Sonnenburg, 2019)—shape oral microbial variation. Several oral microbiome papers have shown “industrialization signatures” in oral microbiomes characterized by declining microbial diversity, enrichment of disease-associated taxa, and compositional convergence across geographically distinct urban populations. For instance, Clemente et al. (2015) observed distinct oral microbial signatures between the Yanomami of Venezuela (an indigenous group with limited Western influence) exhibited higher oral bacterial diversity and greater representation of genera such as *Neisseria* and *Haemophilus* compared to urban U.S. cohorts. (Clemente, et al., 2015). Similarly, *Li et al*. (2014) and *Nasidze et al*. (2009) found that the oral microbiota of rural African and Alaskan Indigenous communities differed substantially from that of industrialized Europeans (Li et al., 2014; Nasidze et al., 2009), while Bisanz *et al*. (2015) observed semi-urban Tanzanian groups displaying intermediate microbial profiles between traditional and industrialized lifestyles (Bisanz et al., 2015). Studies by Lassalle *et al*. (2018), Nasidze *et al*. (2011), and Ryu *et al*. (2024) further identified additional oral microbiome differences across discrete populations with differing diets and subsistence patterns, but often in isolation, without controlling for variation in ecology, history, or host genetics (Lassalle et al., 2018; Nasidze et al., 2011; Ryu et al., 2024). Together, these findings point to declining oral microbial diversity and increased prevalence of disease-associated taxa along gradients of industrialization and dietary homogenization. However, most such studies involved small sample sizes and isolated population comparisons, making it difficult to determine whether observed differences reflect universal processes or context-specific adaptations.

Beyond modern studies, ancient DNA sequencing of dental calculus has begun to reveal how the oral microbiome has transformed alongside major shifts in human subsistence, diet, and health (Adler et al., 2013; Weyrich, et al. 2027; Quagliareillo, 2022). Dietary adaptations during the transition from foraging to agriculture, increased carbohydrate consumption and corresponded with the enrichment of pathogenic taxa such as *Tannerella forsythia* and *Treponema denticola* and a decline in species associated with oral health (Adler et al., 2013; Quagliariello et al., 2022). Specifically, ancient oral microbiome research revealed an increased abundance of potentially pathogenic bacteria (*Porphyromonas gingivalis, Tannerella forsythia, Treponema denticola*, and *Eikenella corrodens)* as Europeans shifted to a more agriculture centric lifestyle during the Neolithic Revolution (Quagliariello et al., 2022). Later industrial transitions further altered community composition, reducing overall diversity and favoring disease-associated taxa (Proctor et al., 2019; Rosier et al., 2014). Recent research using ancient DNA has identified additional microbial shifts linked to the Industrial Revolution, suggesting that “industrialization” continues to influence oral microbial communities in complex and evolving ways (Adler et al., 2013; Quagliariello et al., 2022). However, “industrialization” as a descriptive term does not fully capture the heterogeneity of modernization or its linkages to changing diets, lifestyles, and the uneven global adoption of industrial practices (Benezra, 2020). Some scholars define industrialization more broadly as shifts in subsistence strategies—from foraging to agriculture, or from agrarian to production-based economies—that fundamentally alter human ecology (Inikori, 2002). However, these interpretations remain contested. For example, recent studies have argued that the introduction of farming in southern Europe and other historical events in Europe did not markedly alter oral microbiome composition, suggesting a remarkable degree of continuity across the Neolithic (Ottoni et al., 2021; Fellows-Yates, et al. 2021). This disagreement has fueled a central debate: did the oral microbiome respond to cultural and dietary shifts in similar ways to the gut, such as those associated with industrialization?

However, the totality of industrial mechanisms driving these transformations in the oral cavity remains unclear. For example, adaptations to low-fiber and high-fat diets and the widespread use of antimicrobials explain much of the microbial restructuring observed in the gut (Hamamah et al., 2023; Oliver et al., 2022; Shen et al., 2025; Tuohy & Daniele Del Rio, 2015) but have limited impacts on oral microbiome composition (Belstrøm et al., 2016; Zaura et al., 2015). Others find minimal or no significant differences once technical, environmental, or host-related factors are accounted for. For instance, Belstrøm et al (2014) showed that short-term changes in diet or hygiene had negligible effects on overall oral microbial structure (Belstrøm et al., 2014). Similarly, comparative analyses of modern populations found minimal differences in dental plaque microbiome composition among foragers, agriculturalists, and industrialized groups, further indicating that oral microbial communities are highly conserved across subsistence strategies (Velsko et al., 2023). These inconsistent findings may suggest that previously reported differences primarily reflect variation in sequencing methods, sample preservation, host-related factors, or even the specific oral niche examined. Addressing this uncertainty requires comparative studies that examine oral microbiomes across diverse ecological and cultural settings, using standardized methods that can disentangle the effects of industrialization from those of geography, genetics, and environment.

To examine these patterns in a controlled setting, we focus on a natural experiment in southwestern Uganda, where the Indigenous Batwa (Twa) and their agricultural neighbors, the Bakiga, live in close proximity yet embody contrasting subsistence histories (Nasidze et al., 2011; Scarpa et al., 2021). The Batwa were historically rainforest hunter-gatherers, relying on wild game, honey, and tubers, until their displacement from the Bwindi Impenetrable Forest in 1991 during the creation of the national park (Nasidze et al., 2011; Scarpa et al., 2021). Forced relocation to agricultural settlements has led to profound social and nutritional disruption, including reduced dietary diversity and increased dependence on cultivated crops and aid foods (Disko & Helen, 2014; Ohenjo et al., 2006). In contrast, the Bakiga migrated into the same region approximately 2,000 years ago as part of the Bantu expansion and developed long-standing farming traditions (Pakendorf, Bostoen, and De Filippo 2011; Plumptre et al., 2004). Despite now living under similar climatic and environmental conditions, these two populations possess distinct histories and different timelines of agricultural adoption that allow us to examine how culture and subsistence shape microbial communities. Earlier genetic studies on the population further facilitate our ability to ascertain contributions of genetics into cultural and environmental differences (Perry, 2014).

In this study, we characterize the oral microbiota of Batwa and Bakiga individuals living in the same region, exploring how differences in subsistence and livelihood history structure microbial diversity and composition, in the context of human genetic processes. We then situate these data within a broader global context by integrating them with oral microbiome datasets from non-industrialized and industrialized populations across multiple continents, reprocessed through a standardized bioinformatic pipeline. This approach enables direct comparison across cultural and ecological gradients, allowing us to test whether the patterns observed locally in Uganda align with global trends associated with industrialization. Specifically, we ask: (1) how do shifts in subsistence influence oral microbiome diversity and structure, and (2) to what extent do these differences reflect broader global patterns of microbial change tied to diet and industrialization? By explicitly engaging in the current debate over whether the oral microbiome reflects cultural and dietary history, our study helps clarify when and how industrialization leaves a measurable microbial signal in the oral microbiome.

## MATERIAL AND METHODS

### Batwa and Bakiga dataset

#### Ethics Approval

The saliva samples used in this study were collected with informed consent and approved by the Institutional Review Boards of Makerere University, Uganda (protocol 2009-137) and the University of Chicago, USA (16986A). Approval for this study was also obtained from the Adelaide University, Australia (3808).

#### Sample Collection

The study consists of 100 saliva samples from two Ugandan populations, Indigenous Batwa (n=90) and Bakiga (n=10), randomly selected from a more extensive collection of saliva samples used in a population genetics study into the evolutionary history of the pygmy phenotype in African rainforest hunter-gatherers (Perry et al., 2014). Samples were collected over one field season in 2010 into Oragene collection tubes. Samples of adults from both groups were collected from settlements at eight locations in Kanungu district, Uganda: Buhoma, Byumba, Kebiremu, Kihembe, Kitariro, Mpungu, Mukono, and Nteko. The sample included 56 females and 41 males, with the sex of 3 unknown.

#### DNA extraction, library preparation, and sequencing

Saliva was collected in Oragene DNA collection kits (DNA Genotek Inc., Canada), shipped to the United States, and stored at 80°C following shipment to the United States. A volume of 200 µL of saliva per sample was transferred into new tubes and shipped frozen to the University of Adelaide for further analysis. Cells were lysed from saliva in 2 mL tubes using glass beads vortexed with 470 µL EDTA (0.5M), 30 µL SDS (10%), and 20 µL proteinase K (20 mg/mL). DNA was then extracted from the samples using an in-house silica-based DNA extraction protocol previously described (Brotherton et al. 2013). To account for the use of small volumes of saliva, we modified the total volume of DNA binding buffer to the following: 1.6 mL lysis buffer (1.46 mL Guanidinium DNA binding buffer, 8 µL NaCl (5M), 20.80 µL Triton-X 100, 24.53 µL sterile water, 88.88 µL NaOAc (3M)). Two blank extraction controls were included for every extraction batch of twenty samples (n=10). We amplified the extracted samples using polymerase chain reactions (PCR) with specific barcoded primers targeting a stretch of the V4 region of the 16S ribosomal RNA (rRNA) encoding gene as previously described (Adler et al., 2013; Caporaso et al., 2012a).This reaction used a forward primer 515 F and a reverse primer 806 R; each primer included the appropriate Illumina adapter, an 8-nucleotide index sequence i5 and i7, a 10-nucleotide pad sequence, a 2-nucleotide linker, and the gene-specific primer (Caporaso et al., 2012). We amplified all samples in triplicate and included additional PCR negative controls. Each PCR reaction, performed in triplicate, consisted of 18.05 µL sterile water, 1 µL DNA extract, 0.25 µL of Hi-Fi Taq (ThermoFisher Scientific), 2.5 µL of 10X Hi-Fi Taq (ThermoFisher Scientific), 1 µL MgCl2 (50 mM), 0.2 µL dNTPs (25 mM), and 1 µL each of the forward and reverse primers. We used the following PCR conditions: initial denaturing (95°C, 6 minutes); followed by 37 cycles of denaturation (95°C, 30 seconds), annealing (50°C, 30 seconds), and elongation (72°C, 90 seconds); and final extension (60°C, 10 minutes). After amplification, we pooled the triplicate reactions and visualized the PCR products using electrophoresis on a 2.5% agarose gel. We then quantified the samples using Qubit 2.0 (Life Technologies), pooled them at equimolar concentrations, and purified them using magnetic Ampure beads (Beckman Coulter). We quantified the pooled DNA library using Tapestation 2200 (Agilent) and KAPA SYBR Fast Universal master mix qPCR assay (Geneworks). We performed DNA sequencing using Illumina MiSeq 150 bp paired-end chemistry, generating 2,341,792 paired-end reads. We obtained a mean of 21,289 reads per sample.

#### Data processing

All code for data processing is available https://github.com/microARCHlab/UgandaGlobalMicrobiome_2025/tree/main. Briefly, paired-end sequences were demultiplexed using Illumina bcl2fastq software (v1.8.4) (Aronesty, 2013). The raw sequences were imported into the Quantitative Insights Into Microbial Ecology v2 (QIIME2 2022.8) (Bolyen et al., 2019) using the Earth Microbiome Project protocol, demultiplexed using the qiime2 demux emp-paired feature in QIIME2. Data was trimmed to 150nt for sequence quality control using the quality-filter q-score by denoising with Deblur (Amir et al., 2017). The resultant Amplicon Sequence Variants (ASVs) were aligned with MAFFT (Katoh et al., 2002) to perform a multiple sequence alignment of the sequences. A phylogenetic tree was constructed with the qiime2 align-to-tree-mafft-fasttree pipeline from the q2-phylogeny plugin (Price et al. 2010). We conducted decontam in R (Davis et al., 2018) applying a threshold of 0.60. Taxonomy was assigned using a pre-trained naïve Bayes classifier and the QIIME2 “q2-feature-classifier” plugin. The classifier was trained on the Silva 138 99% reference tree OTUs reference data set (Quast et al., 2013; Bokulich et al., 2018; Robeson et al., 2021). Afterward, ASVs assigned to chloroplast or mitochondria or identified as contaminants in extraction blank controls (EBC) and no template controls (NTC) using decontam were filtered out. Individual “11492.A19489.TW99” and individual “11492.A19451.TW51”, which had a high prevalence of contaminant (46.592% of Synechococcus) and (45.833% of JG30-KF-CM45;g_JG30-KF-CM45) were also filtered out. Sample “11492.A19451.KG59e” was also removed, as it was a duplicate sample. After these, 97 individuals (89 Batwa HG and 8 Bakiga AG) were retained. Features with a frequency below five were also removed from all samples. We decided against filtering out taxa present in less than five individuals from the Ugandan samples because found that we lost some commonly known oral microbes like *Neisseria* and *Provetella*, resulting in a loss of significance in beta diversity. Consequently, we decided not to filter based on sample occurrence. Core diversity metrics were generated using QIIME2 “q2-diversity” (Lozupone et al., 2007; Lozupone & Knight, 2005) with a sampling depth of 4,699. To assess the impacts of unequal sample size, an equal number of Batwa (8) and Bakiga (8) populations were randomly selected, and core diversity metrics were also rerun on this subset of individuals.

We computed alpha and beta diversity using the QIIME2 “q2-diversity” plugin, setting rarefaction at 4,699 sequences per sample(Lozupone et al. 2011; Lozupone and Knight 2005). Alpha diversity for single metadata categories was compared with the Kruskal-Wallis test. The unweighted UniFrac distance between samples was tested with a non-parametric PERMANOVA and ADONIS (Anderson, 2001). Additionally, biplots were included in PCA plots using the Aitchison distance matrix to identify the microbial taxa driving microbial variation across the two groups (DEICODE) (Martino et al., 2019). Differential abundance testing was performed using Multivariable association discovery in population-scale meta-omics studies (MaAsLin2) in R (Mallick et al., 2021).

#### GWAS studies

913,651 SNPs from Illumina HumanOmni1-Quad genotyping array were obtained from publicly available data, corresponding to 82 Batwa and 9 Bakiga individuals in this study. As some individuals in this data set are related, population stratification and family substructure were accounted for using a fixed effect covariate that measures ancestry and a mixed model that includes family structure as a random effect. GEMMA was used to find associations between host SNP profiles and summarized ASVs identified within the salivary data. Covariates included in the analysis included Batwa admixture, self-identified population designation (Batwa or Bakiga), and sex. ASVs were summarized at 5 taxonomic levels: phylum, class, order, family, and genus. CLR and square root transformations were both included in separate analyses. GEMMA was ran on all taxa: SNP pairs in the data set, and BH correction was applied to all results (alpha = 0.15).

#### Data selection and curation for worldwide analyses

To compare the oral microbiome composition of Batwa and Bakiga with those from other worldwide populations, we selected projects with published data from several platforms. Tanzanian data from the European Nucleotide Archive (https://www.ebi.ac.uk/ena/browser/view/PRJEB14941) (Bisanz et al., 2015), Venezuela data from the microbial study management platform QIITA (Clemente et al. 2015), (https://qiita.ucsd.edu/study/description/10052), Nepal (Ryu et al., 2024) and participants from the American Gut Project (AGP) from the project’s ftp web page http://ftp.microbio.me/AmericanGut/20nov2020-demultiplexed-data/ (McDonald et al., et al. 2018) (Table 1) (See https://github.com/microARCHlab/UgandaGlobalMicrobiome_2025/tree/main for detailed description of data wrangling) (Table SI 1 for full metadata). The selected projects included saliva microbiome samples with V4 16S rRNA gene region data using the same primer sets (Caporaso et al., 2012). The dataset, including respective metadata, was filtered for samples from individuals at a single time point, duplicate samples, adults, and those who had not undergone antibiotic or another study-specific treatment/underlying conditions that may have changed the oral microbiome composition using the QIIME2 filter-samples plugin of the feature-table pipeline (Table SI 2). Features in less than five samples and features with a frequency below five across all samples were filtered out. The resulting BIOM tables post filtering were merged with the Ugandan dataset using the QIIME2 feature-table merge plugin. Filtered data was merged with the Ugandan dataset, resulting in 756 individuals. Decontam was then employed on the entire data set using all the controls provided with a threshold of .60. Next, the same analytical analysis described above was performed on the merged and filtered data. Core metrics diversity was computed with a sampling depth of 2,799 ASVs using QIIME2 “q2-diversity” plugin (Lozupone et al., 2007; Lozupone & Knight, 2005), and samples were rarefied to 1,170. This study provides one of the most comprehensive comparisons of over 700 oral microbiome samples spanning Africa, Europe, Australia, and South America.

**Table 1:** List of datasets used in the analyses.

| Project Name | Data Location | No of Samples Used | Citation |
| --- | --- | --- | --- |
| Microbiota at Multiple Body Sites during Pregnancy in a Rural Tanzanian Population and Effects of Moringa-Supplemented | European Nucleotide Archive<br>https://www.ebi.ac.uk/ena/browser/view/PRJEB14941 | 24 | Bisanz et al., 2015 |
| American Gut Project-prep 1164 (industrialized) | The Projects's FTP site<br>http://ftp.microbio.me/AmericanGut/20nov2020-demultiplexed-data/ | 548 | McDonald et al., 2018 |
| The microbiome of uncontacted Amerindians | QIITA<br>https://qiita.ucsd.edu/study/description/10052 | 16 | Clemente et al., 2015 |
| Nepali oral microbiomes reflect a gradient of lifestyles from traditional to industrialized | Sent on request from the first author | 57 | Ryu et al., 2024 |
| Batwa/Bakiga project | In house data | 97 | This study |

#### Community engagement

Following the completion of the oral microbiome analyses, we engaged in a community consultation and results-sharing process in collaboration with members of the Action for Batwa Empowerment Group (ABEG) in Uganda. We considered it important to share these results with community stakeholders before publication and discuss the findings, gather feedback, and identify appropriate approaches for communicating the results to participating communities. Community engagement activities included visits to several Batwa communities, including both original sample collection sites and additional Batwa communities from which samples were not collected. During these engagements, study findings were communicated in accessible, non-technical formats designed for broad community understanding, including oral presentations and visual materials that emphasized images and simplified explanations to meet community needs. Discussions focused on the role of the oral microbiome in human health, the goals of the original saliva-based research, and broad patterns observed in the data. These engagements also created space for community members to share perspectives on changing foodways, oral health practices, environmental transitions, and broader experiences of social and ecological change within their communities. Feedback from these discussions informed our interpretation of the findings and reinforced the importance of situating microbiome research within local historical, cultural, and environmental contexts.

## RESULTS

### Oral microbiome signal was reliably obtained from Ugandan saliva samples

One of the challenges of conducting oral microbiome research from saliva is low endogenous microbiota concentrations compared to host DNA and contamination that can occur from environmental, laboratory, and technical sources, especially those with limited endogenous DNA (Salter et al., 2014; Weiss et al., 2014). Our samples were initially collected in 2010 and extracted in 2017; thus, we first sought to verify the presence of a robust oral microbial signal from our data against negative laboratory controls, EBCs, and NTCs. The alpha and beta diversity of our controls were compared with the biological samples from the Batwa and Bakiga. Alpha diversity, measured using Faith’s Phylogenetic Distance (PD), revealed that biological samples had significantly higher bacterial diversity compared to controls (Kruskal-Wallis, p = 0.00003), as expected. A beta diversity comparison of the microbial composition between samples and controls, using unweighted UniFrac and Aitchsons, showed statistically significant differences between the controls and the samples (PERMANOVA; Controls_samples; unweighted Unifrac; p=0.001; R^2^=0.03; Aitchison; R^2^=0.051; p=0.002). Decontam identified 302 contaminants across all control types, and all of these contaminants were removed from the data set (1409 total features before and 1107 features after filtering). After filtering, the most abundant species in the biological samples were *Streptococcus, Veillonella, Haemophilus, Neisseria, Prevotella, Rothia, Fusobacterium, Prevotella*, and *Gemella*; all ASVs are known to be oral and are present in the Human Oral Microbiome Database (HOMD) database.

### Evolutionary Signatures Still Evident in the Microbiome of Ugandans

Given that Nasidize et al. (2011) reported higher oral microbiome diversity amongst the Batwa compared to other African agricultural populations from different locations, oral microbiome diversity was examined between thes Batwa and neighboring agriculturalists, the Bakiga. While alpha diversity, measured using Faith’s PD, was not statistically significant differences between the Bakiga and Batwa groups (Kruskal-Wallis, p=0.16), the mean level of diversity was slightly higher among the Batwa Hunter-Gatherers (Figure 1A). This was also true when subsampling the data set to equalize sample sizes in each population (p-value = 0.293).

**Figure 1:**
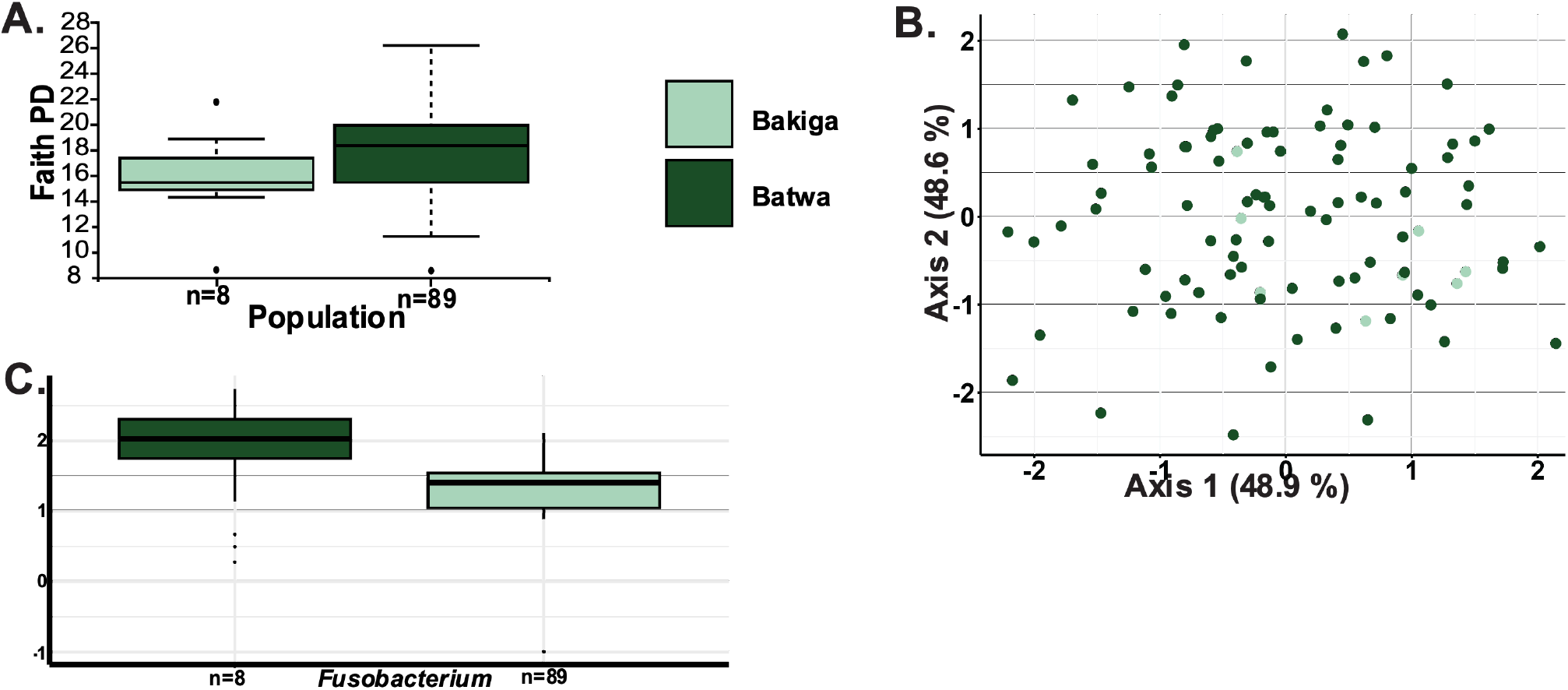
Comparison of alpha, beta diversity, and differential abundance between the Batwa (dark green) and Bakiga (light green) populations in Uganda. (A) Boxplot of Faith’s Phylogenetic Diversity (PD) showing the distribution of alpha diversity values for each population. Boxes indicate the interquartile range, horizontal lines represent the median, and individual points represent samples. (B) Principal Component Analysis (PCA) of microbial community composition based on Aitchison distances. Each point represents a sample and is colored by population. Axes correspond to the first two principal components, which capture the variation in community composition. (C). Results from MaAsLin2 analysis showing the relative abundance of Fusobacterium in each population.

Next, we compared beta diversity across the two populations using unweighted UniFrac and Aitchison distances. While we did not observe apparent population-specific clustering using PCA (Figure 1B), we observed significant differences between the two groups with unweighted UniFrac (PERMANOVA; population; R^2^ = 0.017; p = 0.044) but not with Aitchison distances (PERMANOVA; population; R^2^ =0.022; p = 0.107). These results were also influenced by sample size, as subsampling the Bakiga to create equal sample sizes resulted in significant differences in Aitchison distances (PERMANOVA; population, p = 0.026; R^2^ = 0.21), but not with unweighted UniFrac (PERMANOVA; population, p = 0.189; R^2^ = 0.08). Only one taxa was identified as differentially abundant using MaAsLin2; *Fusobacterium* abundance was significantly higher in the Batwa than in the Bakiga (Figure 1C). Our results suggest that, even though these populations have similar types and amounts of bacteria, the differences in their oral microbiomes revealed by unweighted UniFrac may be linked to evolutionary differences.

### Local geography is associated to Ugandan microbiome differences

Differences in microbiota linked to sex and gender have been observed in the microbiome of other African hunter-gatherer populations (Obregon-Tito et al., 2015; Rosas-Plaza et al., 2022; Schnorr et al., 2014). Thus, we assessed differences within the Batwa and Bakiga populations to identify population-specific factors that may drive diversity within this population. First, we explored the potential for gender-based differences, and no significant differences were observed in alpha diversity between these genders of Batwa individuals (Kruskal-Wallis p =0.6). We then assessed beta diversity using Aitchison’s and unweighted Unifrac and found that gender was significantly different using Aitchison’s distance (PERMANOVA; gender; R^2^ = 0.06; p = 0.019) and not with unweighted Unifrac (PERMANOVA; gender; R^2^ = 0.02; p = 0.492). We explored the differentially abundant taxa between groups but found no significantly differentially abundant taxa between the genders using MaAsLin2.

Next, we assessed diversity, compositional differences, and differentially abundant taxa across the eight settlement sites where the samples were collected. Alpha diversity was significant between the groups (Kruskal-Wallis, p= 0.02), and higher mean diversity in samples collected from Kebiremu (n=11) largely drove this signature (Figure 2). Beta-diversity analysis revealed statistically significant differences in microbial composition across settlement sites (PERMANOVA; collectionsite; Aitchison’s distance: R^2^ = 0.21, p = 0.001; unweighted UniFrac distance: R^2^ = 0.12, p =0.003). Pairwise PERMANOVA comparisons showed that Kebiremu differed significantly from Buhoma (q = 0.036), while other site comparisons were not significant from one another. This difference was again related to differentially abundant levels of *Fusobacterium* across the groups. This suggests that localized factors, such as environmental conditions or unique lifestyle or dietary practices, may shape microbial composition and diversity when sample sizes are sufficient to examine these patterns.

**Figure 2:**
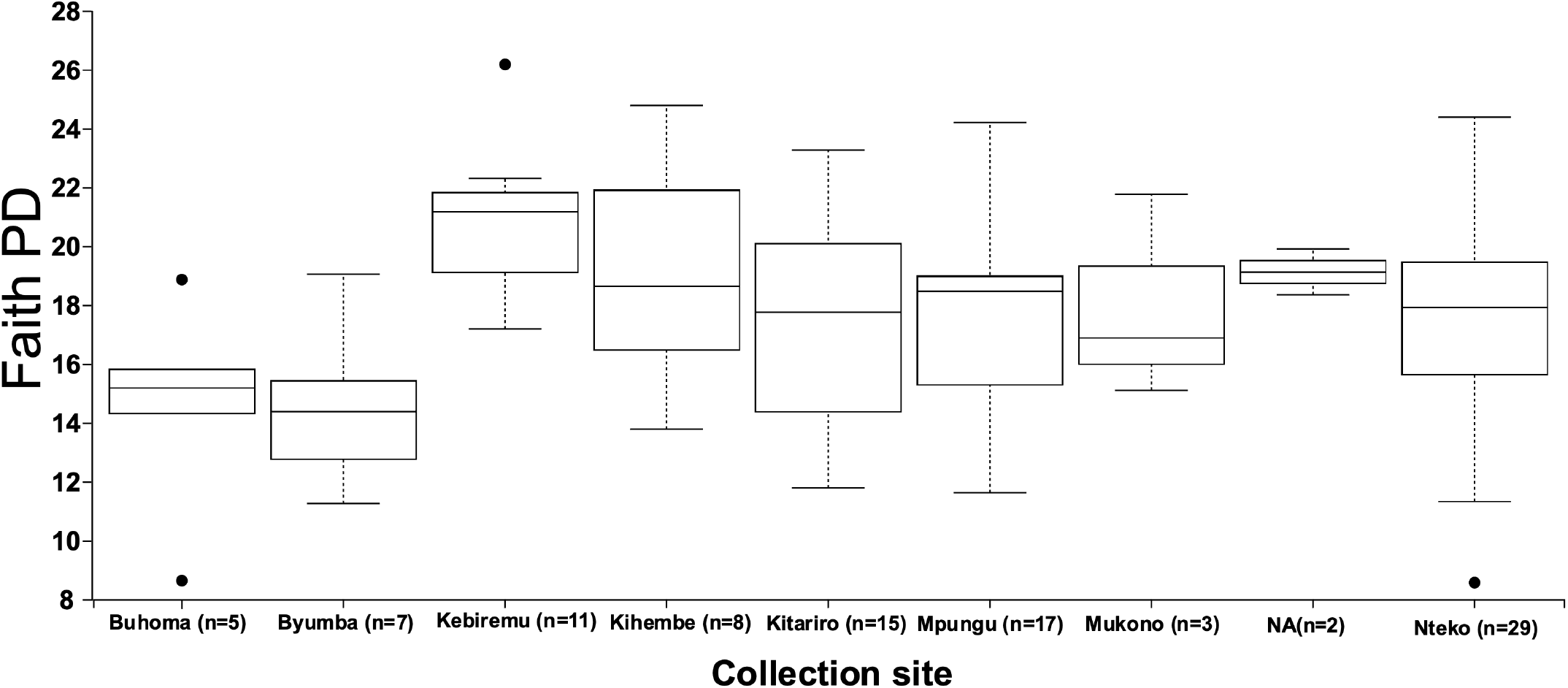
Faith’s Phylogenetic Diversity (PD) across collection sites. Boxplot showing the distribution of alpha diversity values for each collection site.

### Specific SNPs are linked to Fusobacterium presence

GWAS analysis using GEMMA revealed two linkages between specific oral microbes and SNPS within these populations. First, rs7422845(LRP1B) was associated with the presence of *Catonella* in this data set (Figure3A and 3B). This mutation is present in the LRP1B, LDL receptor-related protein 1B, which can be linked to esophageal and oropharyngeal squamous cell carcinoma and periodontitis (Nakagawa et al., 2006). Similarly, increased levels of *Catonella* have been linked to both oral cancers and periodontal disease. Second, an association between rs2938397(PPARG) and *Fusobacteriales* was identified (Figure 3A and 3B). This SNP is within PPARG, which encodes the peroxisome proliferator-activated receptor gamma; enhanced PPAR-gamma expression is linked to the development of Barrett’s esophagus, esophageal adenocarcinomas, and periodontitis in pregnant Japanese women (Hirano et al., 2010a). Similarly, *Fusobacteriales* species can be keystone taxa within the oral microbiome, serving as a species that facilitates large plaque formation necessary for the development of several oral diseases, including periodontitis (Engevik et al., 2021; Han, 2015; Prince et al., 2026), which can influence oral microbiota diversity. Taken together, these results suggest that oral differences across these Ugandan populations may be attributed in part to genetic differences, rather than dietary ones.

**Figure 3:**
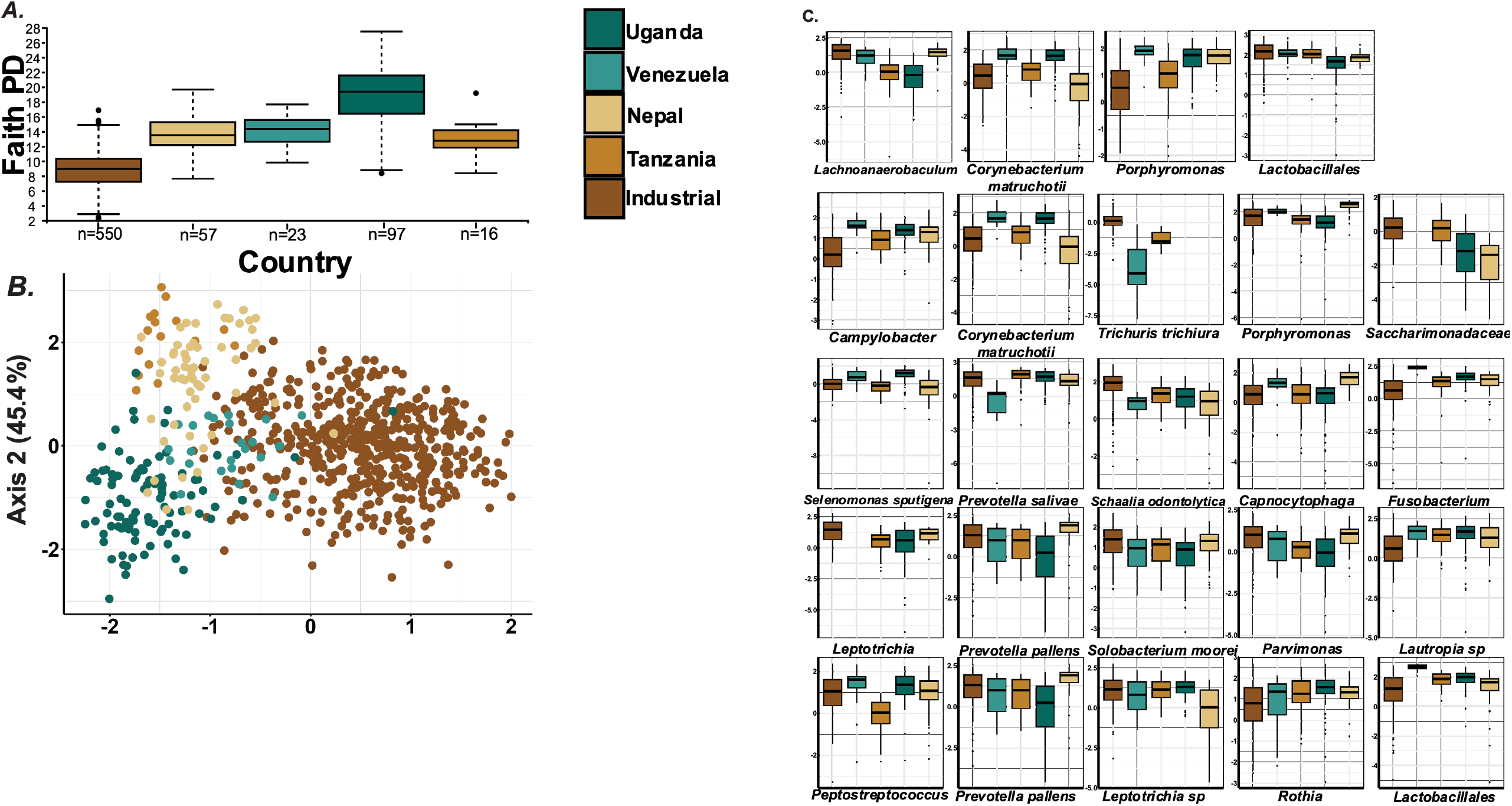
Comparison of alpha diversity, beta diversity, and differential abundance across global populations. A. Faith’s Phylogenetic Diversity (PD) showing variation in alpha diversity among countries. B. Principal Component Analysis (PCA) based on Aitchison distances illustrating beta diversity and clustering of samples by country. C. Differentially abundant taxa identified by MaAsLin2, showing relative abundance patterns of significant taxa across populations.

### Ugandans maintain the highest oral microbial diversity observed to date

Given Ugandan populations were previously observed to have a distinct microbiome compared to global populations, Batwa and Bakiga populations were compared against publicly available V4 16S rRNA data, including that from Tanzanians (Bisanz et al., 2015), Yanomami of Venezuela (Clemente, Pehrsson, Blaser, Sandhu, Gao, Wang, Magris, Hidalgo, Contreras, Noya-Alarcon, et al., 2015), several communities within Nepal (Ryu et al., 2024), and ‘industrialized’ people living within from the United States of America, Europe, and Australia (Flores et al., 2014; McDonald, Hyde, Debelius, Morton, Gunderson, et al., 2018; Ryu et al., 2024). Previous studies have used Greengenes in their taxonomic classification (Clemente, Pehrsson, Blaser, Sandhu, Gao, Wang, Magris, Hidalgo, Contreras, Noya-Alarcón, et al., 2015; Handsley-Davis et al., 2022; Poole et al., 2019). However, we used SILVA to classify our global taxonomic classification, and we found that SILVA was more representative of global diversity by assessing the impact of the different taxonomic databases (Table SI 2). SILVA identified 491 taxa across our dataset, compared to 276 taxa identified using Greengenes (data not shown). Overall, SILVA appeared more representative of the microbial diversity in these populations, so SILVA was employed in all downstream analyses.

Building on this foundation, alpha diversity using Faith PD (Figure 3A) was statistically significantly different across different geographies (Kruskal-Walis; p=6.41 × 10^−78^). The Ugandan population exhibited the highest oral microbial diversity in our global dataset, maintaining 259 ASVs (Table SI 3) primarily from *Actinomyces, Acholeplasma, Capnocytophaga, Cardiobacterium, Treponema, Prevotella, Leptotrichia, Selenomonas*, and *Lentimicrobium* genera. Nine of these ASVs had no hits in the HOMD database (Table SI 4). Significant differences were observed between Ugandans and all other groups, including Uganda and Nepal (q = 2.11 × 10□^13^), Uganda and Tanzania (q = 3.61 × 10□□), and Uganda and Venezuela (q = 3.61 × 10□□) (Table SI 5). However, the largest differences in alpha diversity were observed between industrialized populations, who showed the lowest microbial diversity, and all other populations (Figure 3A); pairwise comparisons using Faith’s PD (Kruskal-Wallis test) identified significant differences compared to Nepal (q = 4.19 × 10□^2^ □), Tanzania (q = 5.48 × 10□^13^), Uganda (q = 6.30 × 10□□^1^), and Venezuela (q = 3.61 × 10□□) (Table SI 5). Overall, this suggests that large quantities of oral microbiome diversity may have been lost in ‘industrial’ countries, similar to observations in the gut. It also suggests that Ugandans and other non-industrial populations maintain higher diversity of oral microbiota than people living in industrial countries.

### Industrialization appears to drive global oral microbiome composition

Composition was assessed across the five population groups using unweighted Unifrac and Aitchison distances matrices (Figure 3B). The industrialized group clustered away from the other groups on Axis 1, with some overlap between Tanzanian samples and industrialized populations (Figure 3B). The clustering indicates regional differences in microbiome composition. Beta diversity significance testing revealed substantial differences in the oral microbiome composition among the five population groups (PERMANOVA; Country; Unweighted UniFrac, R^2^ = 0.20; p = 0.001, Aitchison distances, R^2^ = 0.46; p = 0.001). Pairwise PERMANOVA analyses revealed that microbiota composition in industrialized populations differed significantly from those in Nepal, Tanzania, Uganda, and Venezuela (q = 0.001; Table SI 6). Significant pairwise distinctions were also observed among the non-industrialized groups, (q = 0.001 for all comparisons) (Table SI 5). This observation demonstrates a clear differentiation in microbiome composition between industrialized populations and others.

Using MaAsLin2, we identified 285 taxa that are significantly differentially abundant across countries (Table SI 6). Among these, several taxa show clear population-specific patterns (Figure 3C). For example, *Selenomonas sputigena* was significantly enriched in Ugandan individuals, whereas *Rothia* species were depleted in Uganda but enriched in Venezuelans. Members of *Prevotella*, including *P. salivae* and *P. nigrescens*, showed variable abundance patterns, with enrichment in some populations and depletion in others, reflecting complex population-specific distributions. Additionally, *Lachnoanaerobaculum* displayed contrasting patterns, with higher abundance in Venezuelan and Nepal samples and depleted in Tanzanian and Ugandan individuals. Other notable taxa include *Leptotrichia buccalis*, which is enriched in Nepal but depleted in Venezuela. Overall, these results indicate substantial microbiome variation across populations, with Ugandan individuals showing the most distinct divergence from the Industrial cohort, highlighting the influence of geography, diet, and lifestyle on oral microbial communities.

### Non-Industrial Signatures Are Differentiated by Geography and Subsistence

Given the large effect of industrialization driving global compositional variation, we removed industrial microbiota samples from the dataset and examined diversity within non-industrialized contexts only. Consistent with earlier results, variation in alpha diversity was largely attributed to population group (Kruskal–Wallis; q = 5.84 × 10□^1^ □), with microbiota from Uganda again exhibiting the highest diversity based on Faith’s Phylogenetic Diversity (Figure 4A). Pairwise comparisons revealed significant differences between Nepal and Uganda (q = 1.79 × 10□^12^), Tanzania and Uganda (q = 8.88 × 10□□), and Uganda and Venezuela (q = 4.06 × 10□□). No significant differences were observed between Nepal and Tanzania (q = 0.48), Nepal and Venezuela (q = 0.15), or Tanzania and Venezuela (q = 0.48), suggesting greater similarity in alpha diversity among these non-African populations compared to those within Africa (Table SI 5).

**Figure 4:**
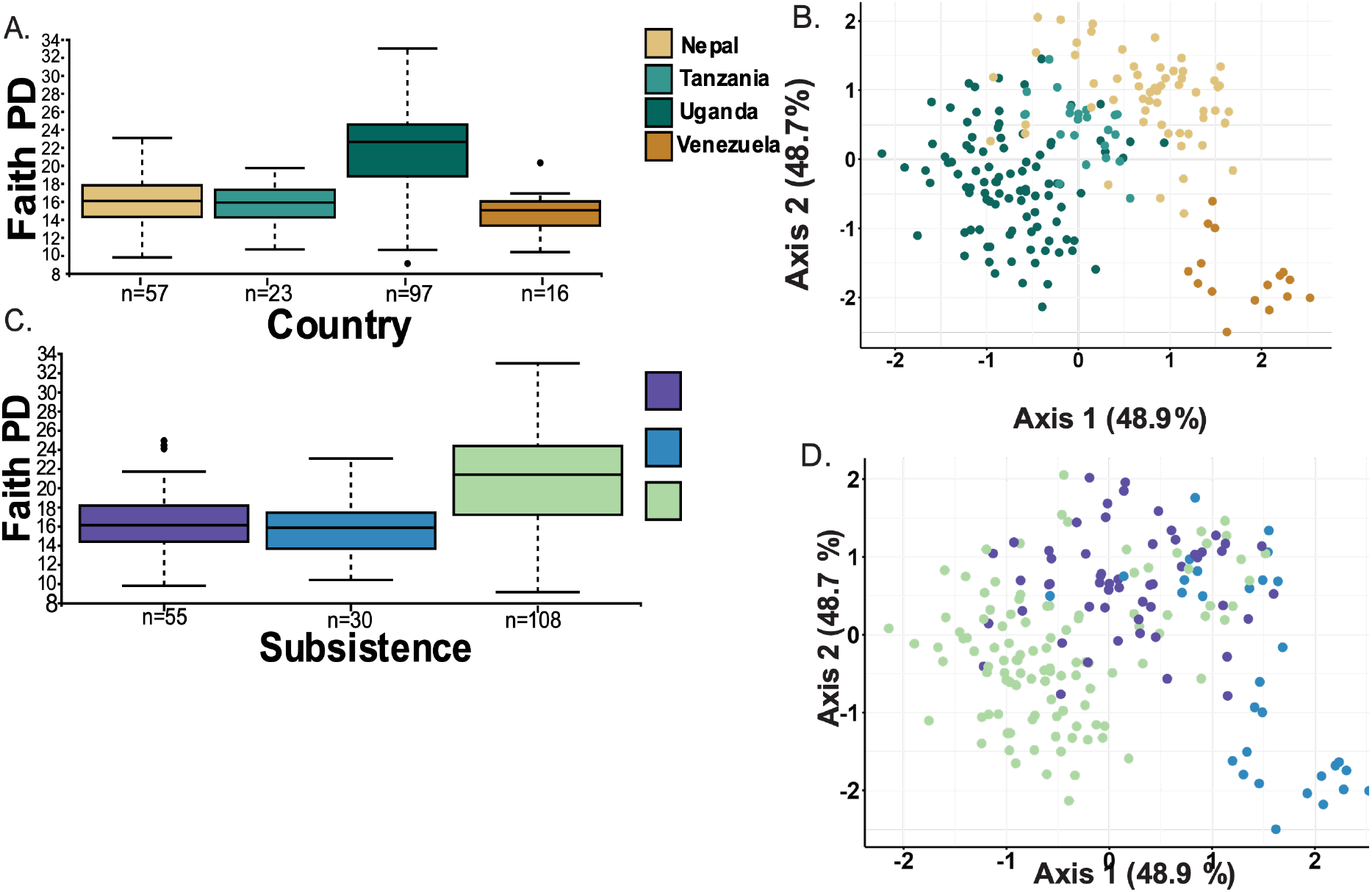
Comparison of alpha and beta diversity among non-industrial populations and across subsistence groups. A, Faith’s Phylogenetic Diversity (PD) showing variation in alpha diversity across non-industrial populations from Nepal, Tanzania, Uganda, and Venezuela. Boxplots indicate Uganda exhibiting the greatest spread in PD values. B, Principal Component Analysis (PCA) based on Aitchison distances showing clear separation of samples by country, reflecting distinct community structures and compositional differences among non-industrial populations. C. Faith’s Phylogenetic Diversity (PD) showing variation in oral microbial alpha diversity among agriculturalist, forager, and transitional groups, with transitional populations exhibiting the greatest alpha diversity. C, PCA based on Aitchison distances illustrating beta diversity patterns. Samples cluster by subsistence strategy, indicating the foragers as being the most compositionally distinct of the group.

Compositional variation using Aitchison distances revealed clear differences linked to African and non-African populations (Figure 4B). Beta diversity analysis based on Aitchison distances showed significant differences in oral microbiome composition among countries (PERMANOVA, q = 0.001). Pairwise comparisons confirmed that all population pairs were significantly distinct (q = 0.001; all comparisons; Table SI 5). To determine whether these differences reflected broader biogeographic structure, we further compared African and non-African populations. Beta diversity analysis using Aitchison distances revealed a strong continental effect (PERMANOVA; q = 0.001), demonstrating that while populations within Africa remain compositionally distinct, they are more similar to one another than to non-African groups Figure 4B). Together, these results indicate that both local (country-level) and continental-scale factors may contribute to global variation in oral microbiome composition.

While continental signatures exist and likely reflect evolutionary, geographic, and environmental differentiation, we further investigated how subsistence strategies can impact these patterns observed in non-industrialized populations. We assigned each person into a subsistence strategy based on data provided by each study (Metadata table, Table SI 0) into one of the four groups: Foragers, Transitional, and Agriculturalist. We appreciate the nuances of subsistence practices in these groups, which are difficult to capture on a global scale. Alpha diversity differed significantly across subsistence groups (q < 0.001; Figure 4C; Table SI 5), which was largely driven by differences between Foragers and Transitional populations (q = 8.29 × 10□□) and between Agriculturists and Transitional groups (q = 5.88 × 10□□), with the Transitional group exhibiting the highest diversity (Figure 4C). However, no significant differences were observed between Agriculturists and Foragers (q = 0.397), suggesting that the process of adapting to a new subsistence strategy may transiently alter alpha diversity before stabilizing (Table SI 6).

Compositional differences across subsistence strategies were also observed using Aitchison distances in a PCA plot (Figure 4D). The Forager group clustered away from the Agriculturist and Transitional groups in a gradient along Axis 1, although distinct clusters between the groups were still present. In this analysis, the Ugandan samples were also divided into two subsistence categories, with the Batwa classified as Transitional and the Bakiga as Agriculturists. Subsistence strategy did explain a significant portion of the variation using Aitchison distances (PERMANOVA, R^2^ = 0.32; p = 0.001) and Unweighted Unifrac (PERMANOVA, Subsistence; R^2^ = 0.10; p = 0.001). Pairwise PERMANOVA analyses identified significant differences in microbiome composition between all pairs of subsistence groups (q = 0.001 for all comparisons; Table SI 4), indicating distinct community structures among Agriculturists, Foragers, and Transitional populations. A comparison of R^2^ values indicates that geographic groupings were able to explain more variance in microbial composition than subsistence strategies (Country: R^2^ = 0.62; Subsistence: R^2^ = 0.32). However, the interaction between country and subsistence was not statistically significant (Pr(>F) = 0.219), suggesting that both geographic location and subsistence strategy are both acting on global oral microbiome diversity. The observed patterns underscore the complex interplay between environmental and lifestyle factors in shaping microbial communities.

## DISCUSSION

In this study, we explored the oral microbiome of the Batwa, a historic hunter-gatherer group, and the Bakiga, an agriculturalist group, both living in the same rainforest environment in Uganda. Based on our results, we propose that a combination of shared environmental, dietary, and genetic factors has likely selected for the similar diversity, but not composition, within oral microbial communities within these two populations. Diversity and composition of microbiota diversity in Uganda compared to other location in the world was driven by both geographic location and their subsistence strategy. Ugandans, similarly, to other non-industrialized locations, maintain higher microbial oral diversity and distinct communities, compared to industrial oral microbiota, suggesting that industrialized practices are a significant driver of oral microbiome diversity, as it is within the gut.

The Batwa are an Indigenous African hunter-gatherer group known for sourcing food through hunting, fishing, and foraging for wild yams and honey in the African rainforest (Disko & Helen, 2014; Ohenjo et al., 2006). In contrast, the Bakiga people migrated into the rainforest during the Bantu expansion and established strong agricultural traditions (Ohenjo et al., 2006). When the Batwa were forcibly displaced from their ancestral lands in 1991 due to the creation of the Bwindi Impenetrable National Park to protect endangered species, their traditional way of life was significantly disrupted. The two groups now share substantial overlap in their oral microbial communities, likely due to shared environmental exposures from the rainforest ecosystem, shared diet, common water resources, and shared disease exposures. Close social interactions, such as the employment of Batwa women on Bakiga farms, may also have facilitated microbial exchange between these populations (Scarpa et al., 2021). These shared microbial taxa could contribute to the lack of distinct culture-specific clustering observed in our beta diversity analyses and the lack of difference in species abundance, suggesting that diet and environmental exposure can also confound evolutionary signatures between populations. Although earlier studies described minimal genetic exchange due to cultural taboos, recent research reveals substantial admixture, likely outside formal marital unions (Patin et al., 2014, 2017; Perry et al., 2014; Perry & Verdu, 2017). These genetic exchanges and shared environmental and dietary exposures may help explain the shared microbial signatures.

Despite these similarities, differences identified between the Batwa and Bakiga in this study are likely attributable to long-standing cultural practices or distinct genetic histories, rather than current environments or dietary differences. We revealed that while *Fusobacterium* was higher in the Batwa, our preliminary GWAS analyses linked this to an association with a specific human genetic variant—rs2938397—located in the PPARG gene (which encodes the PPAR-γ nuclear receptor). This variant is related to disease processes involving mucosal tissues and inflammation, suggesting it could mediate relationships with commensal microbes (Boyer et al., 2018; Hirano et al., 2010b; Wang et al., 2011). This is important to know, as *Fusobacterium* species are known to have direct and indirect negative health implications; *Fusobacterium nucleatum* is indirectly linked to periodontal disease, which can be linked to higher alpha diversity, and directly causes colorectal cancer in some populations (Hajishengallis & Lamont, 2012; Han, 2015; Han & Wang, 2013). Understanding linkages and distinctions between the microbiome and environment, diet, and genetics will be crucial to understanding evolutionary processes that shape oral microbiota diversity on a global scale.

On a global scale, industrialization was identified as the biggest driving factor of global microbial diversity and composition, similar to the gut (Obregon-Tito et al., 2015; Sonnenburg & Sonnenburg, 2019). Previously identified industrial mechanisms in the gut, including dietary shifts toward processed foods and refined sugars, increased urbanization, and widespread adoption of modern healthcare practices (Alt et al., 2022; Ryu et al., 2024; Sonnenburg & Sonnenburg, 2019; Steckel, 1999; Szreter, 2004) may also play key roles here. We identified relationships between geography (country) and subsistence strategy, although mechanistic studies moving forward will need to test this in greater detail. We also observed a decline in oral microbiome diversity in industrial populations compared to those living in rural or remote contexts in other countries. Alterations in oral microbial diversity are linked to disruptions in oral and systemic health, with disease states often characterized by compositional and functional imbalances rather than simple losses of diversity (Hajishengallis & Lamont, 2012; Lamont et al., 2018; Rosier et al., 2018). This reduction in diversity, as shown in the industrial group, is thought to drive the rise in chronic conditions linked to oral health, such as periodontal disease (Hajishengallis et al., 2011; Hajishengallis & Lamont, 2012; Marsh, 2003; Rosier et al., 2018), which is less prevalent in traditional societies (Kriss et al., 2018; Steckel, 1999; Szreter, 2004; Tasnim et al., 2017). For instance, the Tanzanian samples from the semi-urbanized Buswelu area of Mwanza show microbial profiles that overlap with Ugandan and Industrialized populations, suggesting that even limited urbanization can initiate microbial shifts associated with industrial diets (National Bureau of Standard, 2013). The transformation of dietary habits and lifestyle choices in industrialized societies has likely led to microbial shifts that contribute to an increased prevalence of oral diseases. These findings emphasize the importance of studying diverse populations, including industrial, semi-industrial, and non-industrial groups, to grasp the global impact of modernization on microbial diversity and health outcomes.

Our results indicate that while subsistence strategies are important, geographic location was a more powerful determinant of microbiome composition. The more substantial effect of country over subsistence pattern on microbial variation suggests that environmental, genetic, and cultural factors tied to geography may overshadow the influence of diet alone. Furthermore, the lack of significant interactions between these countries and communities’ hints at relatively consistent dietary effects across regions rather than localized interactions with environmental variables. This is particularly relevant when considering how globalization, urbanization, and industrialization might unevenly influence different populations. In contrast, groups that maintain traditional lifeways, such as the Ugandan and Venezuelan populations in our study, offer valuable insights into the resilience of the microbiome. The Ugandans, who retain elements of a hunter-gatherer diet, and the Venezuelans, who remain relatively isolated and adhere to traditional practices, display higher microbial diversity. This contrast highlights how geographic isolation and adherence to ancestral dietary habits can buffer against the homogenizing pressures of modernization. The results indicate that while the Tanzanian populations exhibit a gradient between the Ugandan and more urbanized profiles, their semi-urbanized status may account for this intermediate positioning. The clustering closer to taxa associated with carbohydrate-rich, industrial diets suggests that even limited urbanization can shift microbial diversity. These findings underscore the urgency of examining how traditional, non-industrial lifestyles can help preserve oral microbiome diversity. As more societies undergo rapid industrialization, urbanization, or market integration, understanding these distinctions becomes essential for mitigating the health impacts associated with these processes. This may highlight the potential health benefits of diets and lifestyles that are more aligned with human evolutionary history. By broadening our focus to include these diverse groups, we can better understand the long-term consequences of industrialization on human health and develop strategies to preserve microbial diversity in an increasingly urbanized world.

As with all studies, this project has limitations. To accurately characterize the oral microbiome of these populations, we had to tackle critical technical issues, including sample size disparities, sequencing method, contamination, and taxonomic classification. Regarding sample size disparities between the Bakiga agriculturalists and the Batwa hunter-gatherers, we had notably fewer samples from the Bakiga. Recognizing this limitation, we performed subsampling to assess whether equalizing the sample sizes would significantly impact our results. This approach allowed us to evaluate the robustness of our findings, acknowledging that a larger, more balanced dataset, especially with more Bakiga samples, would enhance the reliability of our conclusions. Second, we were limited to only available data using the V4 region of the 16S rRNA gene, as comparisons across regions can drive technical artifacts (Fadeev et al., 2021; Regueira□Iglesias et al., 2023). This limits our ability to integrate our results with previously published data from other locations and studies, such as that of Nasidze et al., 2011. Third, we assessed contamination within the samples throughout the analysis and used decontam to identify and account for contamination. Because some of the included studies lacked negative controls, we used the negative controls available from other comparative datasets within our study to identify likely contaminants. Taxa overlapping with these contaminants were filtered out, along with mitochondrial and chloroplast sequences, accounting for various types of noise and contamination to understand how cultural and lifestyle changes impact the human oral microbiome. Finally, the commonly used databases, developed primarily from European or Western populations, may introduce biases and underrepresent microbial diversity in non-Western groups, which is also reflected in the unclassified ASVs in the HOMD. Hence, the choice of taxonomic databases plays a critical role when classifying oral microbiomes in diverse populations, particularly those underrepresented in current research. Despite these limitations, our results provide meaningful insights into the oral microbiome differences between these populations.

## CONCLUSION

This study shows the importance of studying diverse populations to capture global microbial diversity and identify factors driving oral microbiota and health differences across populations. Our findings highlight the resilience of the oral microbiome in populations that maintain aspects of their ancestral lifestyles, which are in stark contrast to the homogenizing effects of industrialization observed globally. Including underrepresented groups like the Batwa and Bakiga provides crucial insights into mechanisms that challenge research focused solely in industrialized, WEIRD populations. Such studies are essential for uncovering unique microbial profiles shaped by distinct cultural, dietary, and environmental histories and understanding the totality of how oral microbial communities contribute to health. As global urbanization accelerates, preserving these microbial signatures becomes increasingly essential to understand how modern lifestyles impact oral and systemic health. This work emphasizes the need for more inclusive microbiome research that prioritizes diversity, particularly among Indigenous and traditional communities.

## ACKNOWLEDGEMENTS

We would like to thank the communities that supported this project, first and foremost. We would also like to thank fundings received by the Australian Research Council to complete this project (Award FT180100407 to L.S.W.).

